# sppIDer: a species identification tool to investigate hybrid genomes with high-throughput sequencing

**DOI:** 10.1101/333815

**Authors:** Quinn K. Langdon, David Peris, Brian Kyle, Chris Todd Hittinger

## Abstract

The genomics era has expanded our knowledge about the diversity of the living world, yet harnessing high-throughput sequencing data to investigate alternative evolutionary trajectories, such as hybridization, is still challenging. Here we present sppIDer, a pipeline for the characterization of interspecies hybrids and pure species,that illuminates the complete composition of genomes. sppIDer maps short-read sequencing data to a combination genome built from reference genomes of several species of interest and assesses the genomic contribution and relative ploidy of each parental species, producing a series of colorful graphical outputs ready for publication. As a proof-of-concept, we use the genus *Saccharomyces* to detect and visualize both interspecies hybrids and pure strains, even with missing parental reference genomes. Through simulation, we show that sppIDer is robust to variable reference genome qualities and performs well with low-coverage data. We further demonstrate the power of this approach in plants, animals, and other fungi. sppIDer is robust to many different inputs and provides visually intuitive insight into genome composition that enables the rapid identification of species and their interspecies hybrids. sppIDer exists as a Docker image, which is a reusable, reproducible, transparent, and simple-to-run package that automates the pipeline and installation of the required dependencies (https://github.com/GLBRC/sppIDer).

## Introduction

Interspecies hybrids play a large role in both natural and in industrial settings (Dunn and Sherlock 2008; Soltis et al. 2015; Payseur and Rieseberg 2016; Peris et al. 2017c). However, identification and characterization of the genomic contributions of hybrids can be difficult. High-throughput sequencing can be used to address many of the barriers to identifying and characterizing hybrids. With the influx of sequencing data, the quality and number of reference genomes available is increasing at a rapid pace. Population genomic, ecological diversity, and gene expression projects are underway in many fields. These studies are yielding a high volume of short-read data, but determining the best way to leverage these data can be challenging. A key goal of the modern genomic era is to be able to integrate and synthesize these data to further our understanding of natural diversity (Richards 2017), including addressing key questions about the frequency and genomic identities of hybrid and admixed lineages in the wild.

The number of reference genomes available has rapidly increased, but it is not complete in most clades. To avoid the drawbacks of limited reference genomes, several new phylogenetic approaches have been developed that do not require sequence alignments or whole-genome assemblies, such as phylogeny-building approaches using kmers (Fan et al. 2015), de novo identification of phylogenetically informative regions (Schwartz et al. 2015), and local assemblies of target genes (Allen et al. 2015; Johnson et al. 2016). These methods can accurately reconstruct known and simulated phylogenies of pure lineages. However, these methods have not been tested on hybrid or admixed lineages. As hybrids are the result of an outcrossing event between two independently evolving lineages, their origin is inherently not tree-like. Therefore, placing hybrids on a bifurcating tree will not reflect the topology observed with pure lineages. Placing hybrids on a phylogenetic network is more apt, but it is still untested with alignment-free phylogenetic approaches. Other species identification methods based on local assembly of target genes could lead to erroneous identification, depending on which parent the gene of interest is retained from in the hybrid, or could lead to the assembly of a chimeric gene if the hybrid has retained copies from multiple parents. Therefore, in organisms with alternative evolutionary trajectories, such as hybrids with complex genomes, applying alignment-free phylogenetic methods is difficult and could potentially result in imprecise conclusions.

Other methods to detect interspecies hybrids have been adapted from methods developed for intraspecies diversity, such as F_ST_, STRUCTURE analysis, phylogenetic discordance, linkage disequilibrium, and PCA approaches (Payseur and Rieseberg 2016). There are numerous drawbacks to using these methods to detect interspecies hybrids. For example, most definitions of speciation require the cessation of gene flow and the accumulation of sequence divergence well beyond the levels observed between populations, which are therefore beyond the expectations of most of these approaches. Many of these methods were also developed for diploid obligately outcrossing species, which makes problematic their application to allopolyploids or species that primarily undergo selfing or other forms of inbreeding. Indeed, the basic assumptions of these methods, including gene flow, demographic history, and natural selection, are violated by most interspecies hybrids.

Here we present sppIDer as a novel, assumption-free method that rapidly provides visual and intuitive insight into ancestry genome-wide, which will aid in the discovery and characterization of interspecies hybrids. This method maps short-read data to combination genome, built from available reference genomes chosen by the user. sppIDer allows for the analysis and visualization of the genomic makeup of a single organism of interest, facilitating the rapid discovery of hybrids and individuals with other unique genomic features, such as aneuploidies and introgressions. Therefore, sppIDer is an unbiased method that provides unique and intuitive insights into complex genomic ancestry and regions of differing evolutionary history, which can complement existing methods in the characterization of hybrids.

### New Approaches

Here we describe and make available a user-friendly short-read data analysis pipeline that utilizes existing bioinformatic tools and custom scripts to determine species identity, hybrid status, and chromosomal copy-number variants (CCNVs). Short-reads are mapped to a combination reference genome of multiple species of interest, and the output is parsed for where, how well, and how deeply the reads map across this combination genome. A colorful automated output allows end-users to rapidly and intuitively assess the genomic contribution, either from a single species or multiple species, and relative ploidy of an organism. Figure 1 illustrates the basic workflow in a flow chart of each step. An upstream step creates a combination reference genome, which is a concatenation of reference genomes of interest, before the main pipeline is run. The main pipeline starts with mapping short-read data to this combination reference genome. Then, this output is parsed for percentage and quality of reads that map to each individual reference genome within the combination reference and percentage of unmapped reads; this summary is then plotted so these metrics can be visualized. In parallel, the mapping output is analyzed for depth of coverage. Reads with a mapping quality (MQ) greater than three are retained and sorted into the combination reference genome order; then, coverage across the combination reference genome is computed. A custom script then calculates the mean coverage for each species, and the combination reference genome broken into windows. The output of these analyses is then plotted so that coverage across the combination reference genome can be visualized.

**Figure 1.**
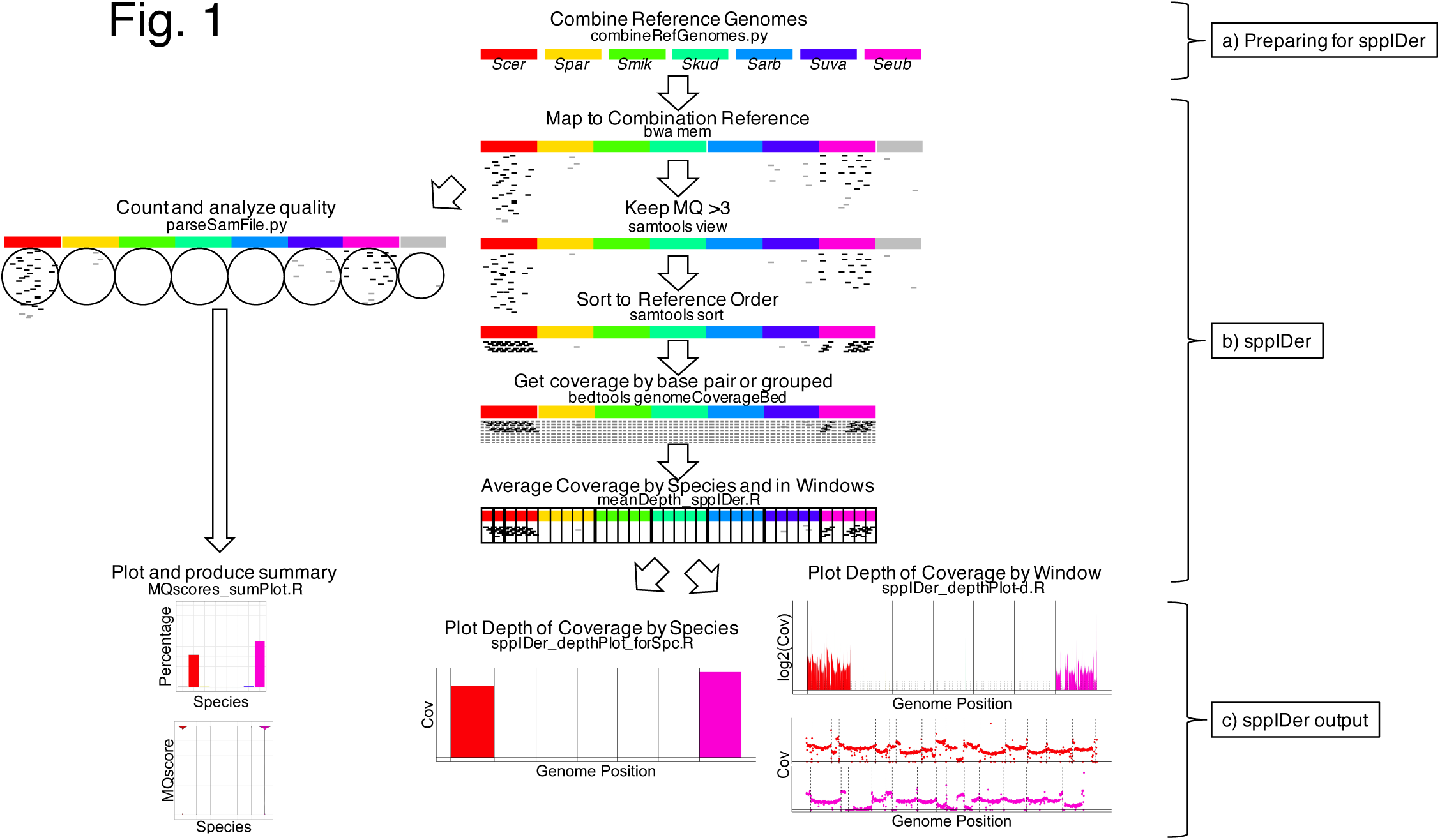
Workflow of sppIDer. (a) An upstream step concatenates all the desired reference genomes (represented by colored bars). Generally, references should be distinct species (see methods for advice about choosing references). This combination reference genome can be used for many analyses. (b) The main sppIDer pipeline. First, reads (short lines) are mapped. This output is used to parse for quality and percentage (left) or for coverage (right). On the left, quality (high MQ black lines versus low MQ light lines) is parsed, and the percentage of reads that map to each genome or do not map (grey bar) is calculated. To determine coverage, only MQ>3 reads (black lines) are kept and sorted into the combination reference genome order. These reads are then counted, either for each base pair or, for large genomes (combination length >4Gb), in groups. Then, the combination reference genome is broken into equally sized pieces, and the average coverage is calculated. (c) Several plots are produced. Shown here are examples of Percentage Mapped and Mapping Quality plots, a plot showing average coverage by species, and two ways to show coverage by windows with species side-by-side or stacked. *Scer* = *S. cerevisiae, Spar* = *S. paradoxus, Smik* = *Saccharomyces mikatae, Skud* = *S. kudriavzevii, Sarb* = *Saccharomyces arboricola, Suva* = *S. uvarum, Seub* = *S. eubayanus*.

We have given this computational pipeline and wrapper a portmanteau of the pluralized abbreviation of species (spp.) and identifier (IDer), to reflect its ability to identify hybrids of multiple species. sppIDer also detects CCNVs, such as those caused by aneuploidy and other genomic changes that do not meet the textbook definition of aneuploidy, including interspecies loss-of-heterozygosity events, interspecies unbalanced translocations, and other differences in relative ploidy. sppIDer is provided as an open source Docker (http://www.docker.com), which organizes the pipeline and all the dependencies into a reusable, reproducible, transparent, and simple-to-run package (https://github.com/GLBRC/sppIDer).

Here we present several applications of sppIDer in yeast, plant, and animal genomes. Through simulations, we show that sppIDer can detect hybrids of closely or distantly related species, and of recent or ancient origin. We use the genus *Saccharomyces* to 1) detect both interspecies hybrids and pure strains; 2) detect hybrids, even with missing reference genomes; and 3) determine how divergent lineages and poor-quality data and reference genomes affect sppIDer’s performance. Next, we test sppIDer’s utility in non-*Saccharomyces* systems: another yeast genus, *Lachancea*; an animal genus, *Drosophila*; and a plant genus, *Arabidopsis*. Finally, we test an extension for non-nuclear DNA using mitochondrial genome data. Overall, sppIDer is robust to many different inputs and can be used across organisms to provide rapid insight into the species identity, hybrid status, and CCNVs of an organism.

## Results and Discussion

### Species and interspecies hybrid identifications

To test sppIDer, we first used the well-studied genus *Saccharomyces* (Hittinger 2013). Seven of the eight species have reference genomes scaffolded at a near-chromosomal level, and there are many interspecies hybrids (Goffeau et al. 1996; Fischer et al. 2000; Dunn and Sherlock 2008; Liti and Carter et al. 2009; Scannell and Zill et al. 2011; Liti et al. 2013; Baker et al. 2015; Naseeb et al. 2017; Peris et al. 2017c). To test sppIDer’s species-level classification ability for a natural isolate, we used the short-read data available for a *Saccharomyces eubayanus* strain isolated in New Zealand (P1C1) (Gayevskiy and Goddard 2016). The reads from this wild *S. eubayanus* strain mapped preferentially to the *S. eubayanus* reference genome (Figure 2a), as seen by normalized coverage only being above zero for the *S. eubayanus* genome. This strain belongs to the same diverse lineage as the reference strain (Peris and Langdon et al. 2016), but as the first isolate from Oceania, these results show that sppIDer can easily classify, to the species level, a divergent wild strain isolated from a novel environment. To test sppIDer’s utility for industrial strains, we used short reads from an ale strain, Fosters O (Gonçalves et al. 2016). This test shows that this brewing strain is a pure species; the *S. cerevisiae* genome is the only genome that had normalized coverage above zero. However, normalized coverage differed within the *S. cerevisiae* genome (Figure 2b & Figure S1a), implying aneuploidies. Coverage was lower for chromosomes VII and XIV and increased for chromosome XIII, in comparison with the genome-wide average coverage, indicating that there are more copies of chromosome XIII and fewer copies of chromosome VIII and XIV. Additionally, we could detect regions of CCNV within a chromosome, such as the small region within chromosome VII where the normalized coverage returned to the genome average.

**Figure 2.**
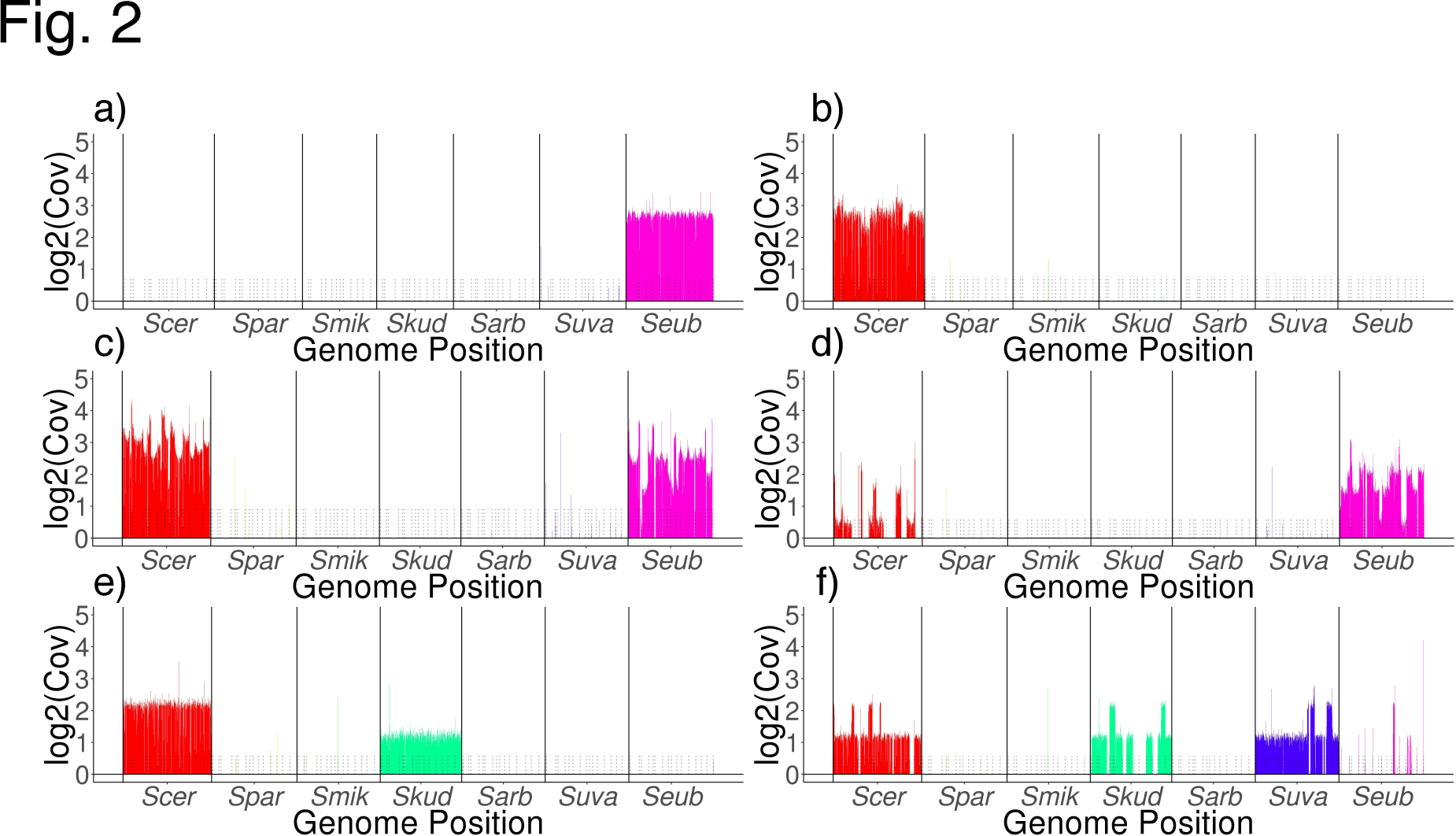
Normalized coverage plots of *Saccharomyces* test cases. (a) Reads from a New Zealand isolate of *S. eubayanus*, P1C1, mapped to the *S. eubayanus* reference genome (magenta). (b) Reads from an ale strain, FostersO, mapped to the *S. cerevisiae* reference genome (red), with visually detectable aneuploidies. (c) Reads from a hybrid Frohberg lager strain, W34/70, mapped to both the *S. cerevisiae* and *S. eubayanus* reference genomes in an average approximately 1:1 ratio with visually detectable translocations and aneuploidies. (d) Reads from a hybrid Saaz lager strain, CBS1503, mapped to both *S. cerevisiae* and *S. eubayanus* reference genomes in an average approximately 1:2 (respectively) ratio with visually detectable translocations and aneuploidies. (e) Reads from a wine hybrid strain, Vin7, mapped to *S. cerevisiae* and *S. kudriavzevii* (green) reference genomes in an average approximately 2:1 (respectively) ratio. (f) Reads from a hybrid cider-producing strain, CBS2834, mapped to four reference genomes: *S. cerevisiae, S. kudriavzevii, S. uvarum* (purple), and *S. eubayanus*.

To test sppIDer’s ability to delineate hybrids, we used short-read data from two *S. cerevisiae* X *S. eubayanus* lager yeast lineages, Saaz (strain CBS1503) and Frohberg (strain W34/70). These results recapitulated the known relative ploidy and rearrangements, where ploidy differs both within and between genomes. Specifically, the Frohberg lineage contains approximately two copies of each chromosome from both *S. cerevisiae* and *S. eubayanus*. Thus, what was observed matched this expectation, where the average normalized coverage across both the *S. cerevisiae* and *S. eubayanus* genomes were approximately at the same level, but there were clear fluctuations, indicating ploidy changes (Figure 2c & Figure S1b). In our test with a representative of the Saaz lineage, we observed that the *S. cerevisiae* genome had an average normalized coverage of ∼0.5, that fluctuated from none to two, and the *S. eubayanus* genome had an average normalized coverage of 1.5, that fluctuated from zero to three (Figure 2d & Figure S1c). These results match with previous observations that the Saaz lineage is approximately haploid for the *S. cerevisiae* genome and diploid for the *S. eubayanus* genome. Additionally, from the sppIDer plots, we also easily inferred the previously described aneuploidies and translocations (Figure 2c-d) (Dunn and Sherlock 2008; Okuno et al. 2016).

As an additional hybrid test, we used short-read data from the wine strain Vin7, a *S. cerevisiae* X *Saccharomyces kudriavzevii* hybrid. From the normalized coverage plot (Figure 2e), we could determine that Vin7 has retained complete copies of both parental genomes, but at different ploidy levels. Specifically, the normalized coverage for *S. cerevisiae* was around two across the genome, while the normalized coverage for *S. kudriavzevii* was consistently around one across the genome. Here we could infer that this strain has double the number of copies of *S. cerevisiae* chromosomes as it does of *S. kudriavzevii* chromosomes. Although exact ploidy cannot be measured without direct measures of DNA content, the inferred ploidy is consistent with previous studies (Borneman et al. 2012; Peris et al. 2012; Borneman et al. 2016).

As a final test of interspecies hybrids, we used data from the cider strain CBS2834 (Almeida et al. 2014). Here sppIDer detected large genetic contributions from *S. cerevisiae, S. kudriavzevii*, and *Saccharomyces uvarum,* as well as introgressed contributions from *S. eubayanus* (Figure 2f & Figure S1d). Although the *S. eubayanus* genetic contribution is quite small, seen on chromosomes XII and XIV, it was still easily detected by sppIDer. These examples show that sppIDer can easily detect higher-order interspecies hybrids, even those with minor contributions from several species.

### Testing the limits of sppIDer with a simulated phylogeny

To test sppIDer’s performance with hybrids of varying levels of parental divergence, we used a simulated phylogeny. To build this phylogeny we started with the *S. cerevisiae* reference genome and produced a phylogeny of 10 species through several rounds of simulating short-read sequencing data, applying a set mutation rate, and assembling those reads. For these simulated genomes, sister species were ∼4% diverged, and the most distantly related species were ∼20% diverged (Figure 3a). This simulated phylogeny allowed us to test pseudo-hybrids from closely and distantly related lineages. Further, the iterative process of phylogeny building allowed us to create ancient pseudo-hybrids that simulated the result from hybridization of a common ancestor predating a lineage split. sppIDer accurately mapped pure lineages to their corresponding reference genome (Figure 3b). For all 10 species, > 90% of the reads mapped to their corresponding reference genome. The read simulation and assembly process resulted in varying quality final references, but despite differences in genome quality, all reads still mapped accurately and were not biased to the best reference genome.

To determine sppIDer’s applicability to hybrids of both closely and distantly related parents and of recent and ancient origin, we tested sppIDer with pseudo-hybrids of different combinations of simulated species. sppIDer accurately detected all true hybrid parents. When pseudo-hybrids were between sister species, <0.01% of the reads mapped promiscuously to other species (Figure 3c). When we used more divergent pseudo-hybrids, sppIDer still detected the true parents, with <5%of the reads mapped promiscuously to the sister species (Figure 3d). Additionally, we simulated ancient pseudo-hybrids, between common ancestors before lineage splits, and found that sppIDer mapped the reads of these hybrids to the references of the lineages that descended from the ancestors that hybridized (Figure 3e). With complete knowledge of this simulated phylogeny, we were able to test many different potential hybrid arrangements and found that sppIDer detected the true parents of all hybrids.

**Figure 3.**
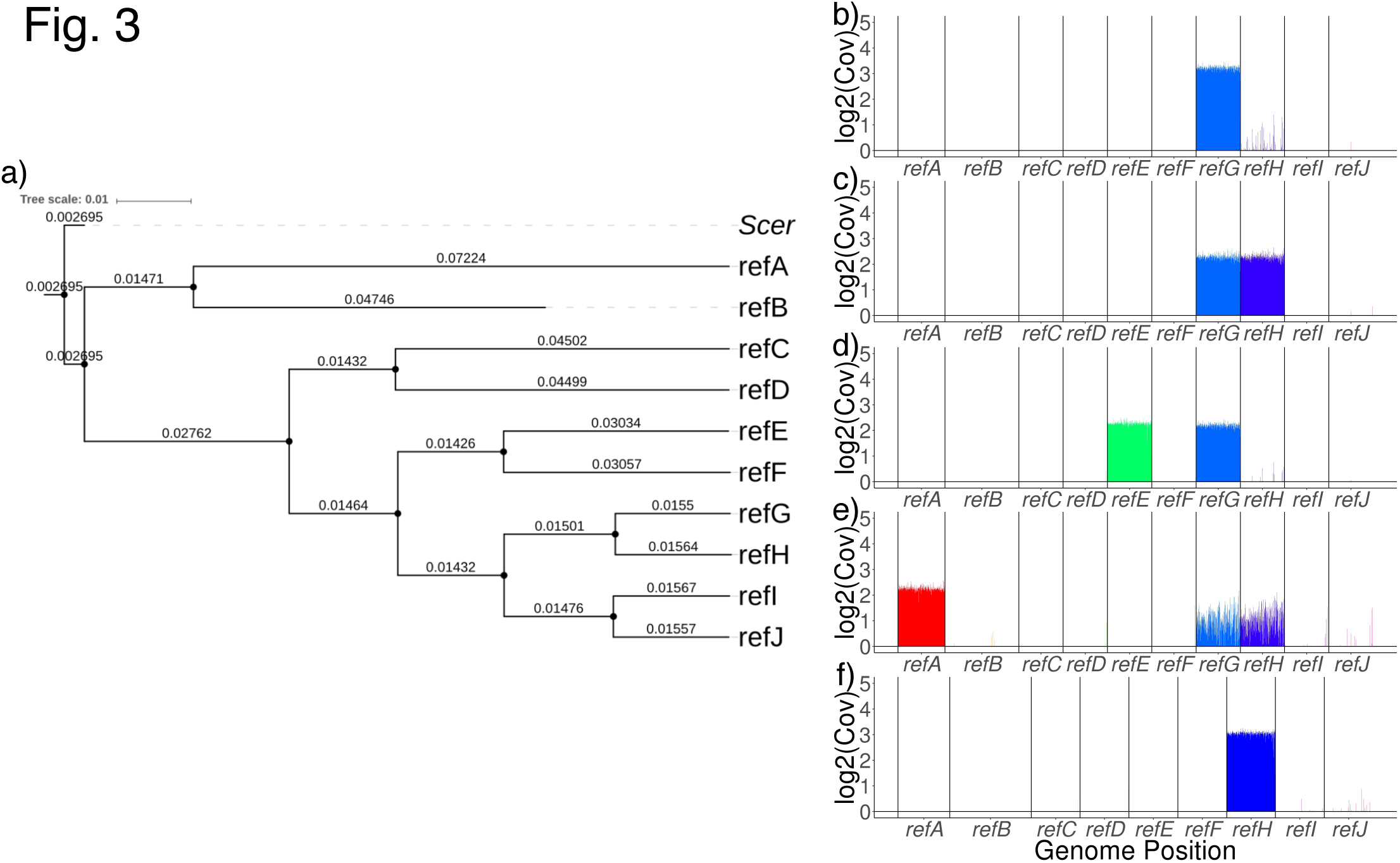
Simulated phylogeny of 10 species and sppIDer’s detection of hybrids from this phylogeny. (a) Phylogeny built with AAF. (b) Reads from G mapped to the G reference genome. (c) Reads from a pseudo-hybrid of the closely related species G and H mapped to the G and H references. (d) Reads from more distant pseudo-hybrid of E and G mapped to references E and G. (e) Reads of ancient pseudo-hybrid of A and a common ancestor of G and H mapped to the references of A, G, and H, which are the lineages that descended from the hybrid’s parents. (f) Without the G reference genome, reads from a pseudo-hybrid of the closely related species G and H mapped to the H reference genome, with some mapped promiscuously to references I and J.

Finally, we tested a scenario, which is common in biology, of incomplete knowledge of the clade of interest. This dearth could due to many variables, such as a described species lacking a reference genome or a species being unknown to science altogether. To test the effect of missing a species, we removed one species’ reference genome from the combination reference genome, then mapped pure lineage and pseudo-hybrid reads to this permuted genome. With reads of a simulated pseudo-hybrid of sister species, G and H, we observed that, when one parent genome was missing, the reads mapped primarily to the reference genome of the remaining parent, reference H, with slightly increased promiscuous mapping of reads to the next-closest clade, references I and J (Figure 3f). Therefore, with incomplete reference genome knowledge, detecting hybrids of closely related species is limited. However, we could still detect hybrids of more distantly related species, such as a pseudo-hybrid of E and G and a pseudo-hybrid of the common ancestor of A and the common ancestor of G and H (Figure S2), though our inference of parentage was biased by the availability of reference genomes. Therefore, with incomplete knowledge of reference genomes, hybrid detection is limited, and the inference of true parentage can suffer in specific cases, but generally, distant and ancient hybrids can be detected.

### Hybrid detection with missing reference genomes

To empirically address how sppIDer would be affected by missing reference genomes, such as for hybrids whose parents are themselves unknown (Hoot et al. 2004; Pryszcz et al.2014), we focused again on the genus *Saccharomyces*. Specifically, we used the *S. cerevisiae* X *S. kudriavzevii* hybrid (Vin7) and the *S. cerevisiae* X *S. eubayanus* Frohberg lager yeast (W34/70) as examples. We tested the performance of sppIDer on short-read data from both hybrids by removing the *S. cerevisiae* reference genome and, in a separate test, removing the reference genome of the other parent. Our expectation was that reads would map to the genome of the sister species, if it were available, or that they would fail to map or be distributed across other genomes, if there were no close relatives.

When we removed the *S. eubayanus* reference genome for the lager example, the proportion of reads that failed to map increased, as did those reads that mapped to *S. uvarum*, its sister species (∼93%identical in DNA sequence, Libkind and Hittinger et al. 2011), albeit with a decreased mapping quality (MQ) (Figure 4c). We then tested sppIDer on Vin7 and W34/70 when the *S. cerevisiae* reference genome was removed (Figure 4a & d). In both examples, the proportion of reads that mapped to *Saccharomyces paradoxus, S. cerevisiae’s* sister species (∼87% identical in DNA sequence), increased (Figure 4a & d). Thus, the absence of a reference genome for one of the parents of a hybrid led to increased mapping to its sister species, instead. We also tested removing the *S. kudriavzevii* reference genome for Vin7. Since there is not a sister species closely related to *S. kudriavzevii*, the number of unmapped reads increased, and the remaining reads mapped to the reference genomes other species of the genus in approximately equal proportions (Figure 4f).

**Figure 4.**
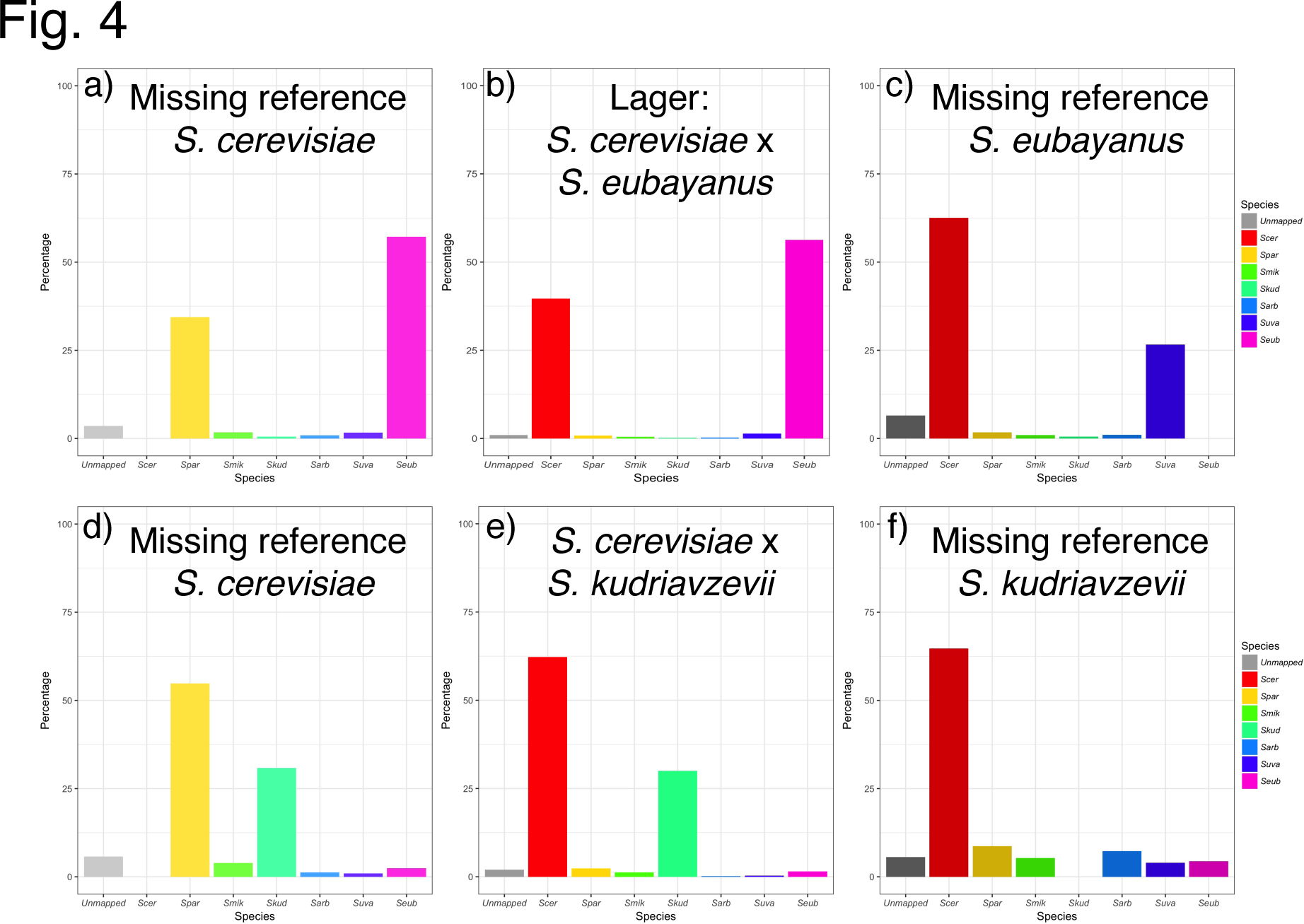
Comparison of the percentage of reads that mapped when different reference genomes were excluded, compared to when all possible reference genomes for *Saccharomyces* were available (middle panels). (a) When the *S. cerevisiae* reference genome was not provided and reads from a Frohberg lager strain, W34/70, were mapped, more reads failed to map (grey) or mapped to the *S. paradoxus* reference genome (yellow). (b) When the full array of *Saccharomyces* genomes was provided, reads for the lager strain mapped to both *S. cerevisiae* and *S. eubayanus*. (c) When the *S. eubayanus* reference genome was removed, more reads from the lager strain failed to map or mapped to the *S. uvarum* reference genome (purple). (d) With the removal of the *S. cerevisiae* reference genome, reads from the *S. cerevisiae* X *S. kudriavzevii* hybrid strain Vin7, which would normally map to *S. cerevisiae*, instead failed to map or mapped to *S. paradoxus*. (e) When all genomes were used, reads mapped to both *S. cerevisiae* and *S. kudriavzevii*. (f) With the removal of the *S. kudriavzevii* reference genome, reads that would normally map to *S. kudriavzevii* instead failed to map or were distributed across all other genomes.

From these tests, we would have easily inferred that W34/70 was a hybrid, regardless of whether either parent genome was withheld (the actual state of affairs for *S. eubayanus* before Libkind and Hittinger et al. 2011). Using the coverage plots, we were still even able to infer the same CCNVs for W34/70 that we observed with the full suite of reference genomes. With Vin7, we still easily inferred its hybrid status without including the *S. cerevisiae* genome. Without the *S. kudriavzevii* reference genome, Vin7 produced an unusually high number of unmapped reads without a decrease in mapping quality to *S. cerevisiae*, a result that should spur the investigator to perform more detailed analyses to search for evidence of contributions by an unknown species, such as de novo genome assembly and phylogenetics. Therefore, even without a full complement of reference genomes, sppIDer can still be useful for rapid inference of interspecies hybrids.

### Hybrid detection with simulated low-quality reference genomes

To test a scenario where not all of the reference genomes are ideal, we used iWGS (Zhou et al. 2016) to independently simulate reads and then assemble de novo genomes for *S. cerevisiae, S. kudriavzevii, S. uvarum*, and *S. eubayanus*. These simulations resulted in reference genomes with many more scaffolds and with a lower N50 than the published genomes (Table S1). These low-quality genomes were independently swapped for the high-quality references in the combination reference genome and tested with short-read data. We started by testing simulated pseudo-lager short reads where we expected reads to map both to the *S. cerevisiae* and *S. eubayanus* reference genomes. Whether we swapped in the low-quality *S. cerevisiae* reference (Figure S3a) or the low-quality *S. eubayanus* reference (Figure S3b), the reads still mapped equally to the references that were used to simulate the reads with minimal promiscuously mapped reads to their sister species reference genomes.

We next tested the limits of sppIDer with the empirical data for CBS2834 because it has the most complex arrangement of contributions from four species. Tests with each simulated low-quality reference genome independently showed that we could indeed recapitulate the same inference of ancestry and that roughly the same proportion of reads mapped to each reference genome (Figure S4) as with high-quality reference genomes (Figure S1d). Here, the inference of approximate ploidy became more difficult, and visually interpreting translocations between species was impossible. When both high-quality *S. cerevisiae* and *S. kudriavzevii* reference genomes were used, we could infer translocations between these two genomes on chromosomes IV, X, and XV due to mid-chromosome ploidy changes that are compensated for in the other genome. There were more promiscuously mapped reads to the high-quality reference genomes of the sister species, but not at the same level as mapped to the true parent reference genomes.These tests with simulated low-quality de novo genomes showed that, both with simulated and empirical data, proper hybrid genome contributions can still be identified, and ploidy shifts still detected, despite the poor-quality reference genomes, but the inference of translocations and ploidies of specific chromosomes becomes difficult.

### Hybrid detection with low-coverage and long-read data

To further explore the power of sppIDer, we wanted to test how little coverage was needed to still detect the proper ancestry (Figure S5). Using data simulated at varying coverages, we found that only 0.5X coverage was needed to recover the true ancestry for a single species (Figure S5a-b), single species with aneuploidies (Figure S5c-d), and interspecies hybrids (Figure S5e-f). We also tested empirical data by down-sampling the FASTQ files of CBS2834 and found that we could still detect contributions from the four species at as low as ∼0.05X coverage (Figure S5g), but we lost the ability to infer ploidy at around ∼0.5X coverage (Figure S5h).These low coverage tests show how powerful sppIDer is, even with scant data, which could be a boon in many systems with large genomes or when sequencing resources are limited.

We also tested sppIDer with simulated PacBio long-read data from the *S. cerevisiae* genome and a hybrid pseudo-lager genome with equal contributions from the *S. cerevisiae* and *S. eubayanus* reference genomes. We found that we could still easily determine the species contribution for each (Figure S6), suggesting sppIDer’s utility will continue if long-read technologies eventually supplant short-read sequencing technologies.

### Divergent lineages and poor-quality data

Since sppIDer relies on reference genomes, we recognized that it might be biased in its ability to work with lineages that were highly divergent from the reference genome, as might be the case in many systems. We tested this scenario with an example from *S. paradoxus*, one of the most diverse *Saccharomyces* species (Liti and Carter et al. 2009; Leducq et al. 2016). Compared to a representative of the reference genome’s lineage, fewer reads from the divergent lineage (∼96%identical) mapped and with poorer quality (Figure S7a-b). We also tested this effect in *S. kudriavzevii* using poor-quality data (36-bp reads from a first-generation Illumina Genome Analyzer run by Hittinger et al. 2010) and found qualitatively similar results, but many more unmapped reads. Thus, while divergence from the reference genome affected map-ability, sppIDer still worked generally as expected. However, when mapping percentage and quality decline substantially, such as seen in these test cases, sppIDer can provide an early indication that the organism may be highly divergent from the reference genome, which may merit further investigation.

### Comparison to alignment-free phylogenetic methods

Alignment and assembly (AA)-free phylogeny-building methods are gaining popularity, but they have not previously been applied to hybrid data. Therefore, we also tested how AA-free phylogenetic methods, such as AAF (Fan et al. 2015) or SISRS (Schwartz et al. 2015), performed in detecting and visualizing hybrids compared to sppIDer. We found that these methods performed well when given only pure lineages, but when hybrids were included, they either failed completely or produced incorrect phylogenies. We tested both our simulated phylogeny and empirical *Saccharomyces* data. For the simulated data, both AAF and SISRS produced the correct phylogeny when given the 10 simulated species. However, when given any hybrid data, AAF failed to produce the correct phylogeny and instead clustered the hybrid with its parents, while SISRS failed to complete at all. With the empirical data, we saw similar results; with AAF, we could recapitulate the phylogeny of the genus *Saccharomyces* when using only pure samples, but when we included any hybrid, an incorrect phylogeny was produced (Figure S8a and c). SISRS had similar issues with producing the correct phylogeny with hybrids, but its output allowed for more nuanced network visualizations. For CBS2834, the SISRS output allowed us to infer the shared background with *S. cerevisiae, S. kudriavzevii, S. uvarum,* and *S. eubayanus* (Figure S8d), but the proportion of contribution from each species was difficult to estimate compared to the sppIDer output. Overall, we found that these methods have serious limitations when used with hybrids, but they could be used as a complement to sppIDer to make inferences about pure parental lineages.

Methods that assemble targeted genes from short read-data, such as aTRAM and HybPiper, can be used with poor-quality references and/or references that may be missing genes of interest. We tested these tools with a panel of loci that can be used to delineate the *S. eubayanus* populations (Peris and Langdon et al. 2016). HybPiper and aTRAM were able to match short-reads to a locus of interest 59%or 34% of the time, respectively, but they could only assemble these reads 23% or 28% of the time, respectively. Neither method could assemble one locus for all 15 strains tested, including both hybrids and non-hybrids (Table S1). While these methods can be powerful when applied in a targeted manner to pure strains, they fail when applied to hybrid data.

### Non- *Saccharomyces* examples

*Lachancea*: refining the interpretation of voucher specimens

With the publication of 10 high-quality *Lachancea* genome sequences (Vakirlis et al. 2016) and another two recently-described and fully-sequenced species (González et al. 2013; Freel et al. 2015; Sarilar et al. 2015; Freel et al. 2016), this genus is becoming a powerful yeast model. As molecular techniques improve, initial identifications in culture and museum collections can yield new interpretations. For example, the strain CBS6924 was initially identified as *Lachancea thermotolerans*, but recent evidence suggested it as a candidate for a novel species (*Lachancea fantastica* nom. nud. Vakirlis et al. 2016). Its closest relative, *Lachancea lanzarotensis,* was also recently described (González et al. 2013). To test sppIDer’s utility for determining whether a strain or voucher specimen is or is not properly classified, we tested mapping reads from CBS6924 to a combination genome with all *Lachancea* reference genomes (Figure S9a), then removing the ‘*L. fantastica*’ reference genome (Figure S9b), and then removing both the ‘*L. fantastica*’ and *L. lanzarotensis* reference genomes (Figure S9c). When both reference genomes were removed, the reads were spread across many genomes, and the initial classification of the strain as *L. thermotolerans* would have been easily falsified. By including the *L. lanzarotensis* reference genome, most reads mapped to that reference, but still poorly enough to warrant additional investigation. When the ‘*L. fantastica*’ reference genome was included, CBS6924 reads mapped unambiguously to this reference. These results demonstrate sppIDer’s utility outside of the genus *Saccharomyces* to aid in reclassifying provisional species identifications of voucher specimens from culture and museum collections.

### Drosophila

To test sppIDer with larger genomes, we examined the animal genus *Drosophila*, which is a large genus with many available reference genomes (Adams et al. 2000; *Drosophila* 12 Genomes Consortium 2007; Alekseyenko et al. 2013; Sanchez-Flores et al. 2016), as well as several species still lacking reference genomes. The difference in reference genome qualities led us to remove contigs <10Kb from our combination reference genome; this paring down sped up computation time, reduced memory usage, and improved the visualization, but otherwise did not affect the results (Figure S10a). To test the ability of sppIDer to distinguish closely related species, we started with the *Drosophila yakuba* species complex (Turissini et al. 2015), where *D. yakuba* has a sequenced reference genome available, but its close relative *Drosophila santomae* and more distant relative *Drosophila teissieri* do not. Here we observed that short reads from a *D. yakuba* (Comeault et al. 2016) representative mapped well to the *D. yakuba* reference genome. As we moved from the close relative *D. santomae* (Figure 5b) to a more distant one, *D. teissieri* (Figure 5c), the mapping percentage and quality decreased with increased promiscuous mapping to other relatives (Figure 5a-c). Thus, as in yeasts, sppIDer can classify pure species and their close relatives well and provide insight to guide downstream analyses.

**Figure 5.**
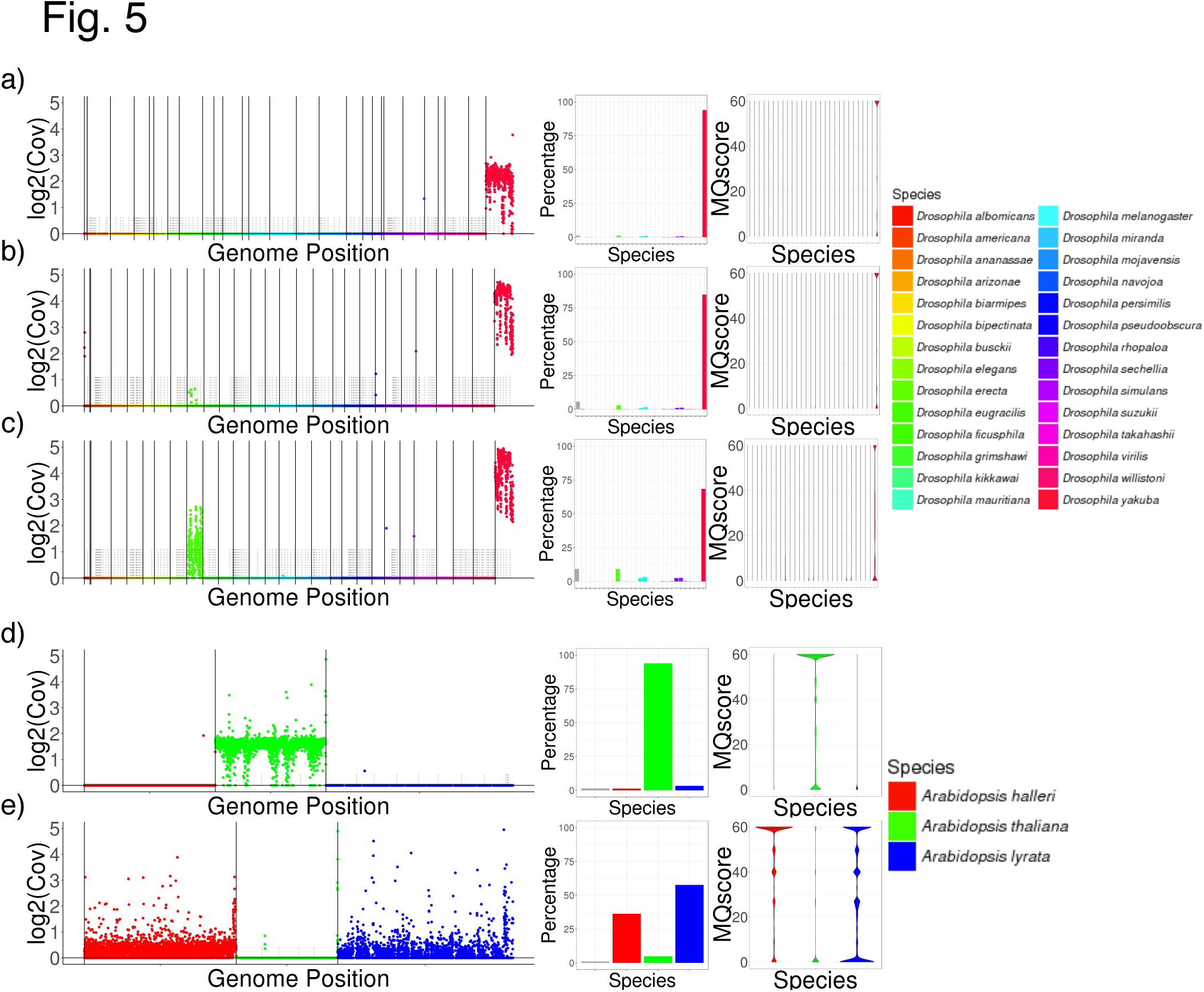
Examples using animal and plant genomes. (a) Reads from a *D. yakuba* individual mapped primarily (> 99%) to the *D. yakuba* reference genome. (b) Reads from the sister species *D. santomae* mapped best to the *D. yakuba* reference genome with some mapped promiscuously to other reference genomes. (c) Reads from the more distantly related species *D. teissieri* mapped mostly to the *D. yakuba* reference genome, but with more reads not mapped and mapped promiscuously to other related reference genomes. (d) Reads from an *Arabidopsis thaliana* accession from Tanzania mapped back to the European reference genome for *A. thaliana*. The repetitive nature of centromeres causes the coverage to fluctuate around those regions. (e) Reads from the hybrid species *A. kamchatica* mapped to the two parental reference genomes: *A. halleri* and *A. lyrata*.

We also used *Drosophila* short-read data to test sppIDer’s ability to detect hybrids in non-fungal systems. In this case, we used genomic data from a pure parent and RNA-seq data from a F_1_ interspecies hybrid (Coolon et al. 2014). We found that sppIDer could easily detect hybrids in an animal model, but as expected, detection of CCNVs using RNA-seq was not possible (Figure S11).

### Arabidopsis

The study of hybrid speciation and allopolyploidy in plants has a long history (Rieseberg 1997; Soltis et al. 2015), and we choose *Arabidopsis* as our plant test case because it has reference genomes available for *Arabidopsis halleri, Arabidopsis thaliana*, and *Arabidopsis lyrata* (Swarbreck et al. 2008). There are drastic differences in the quality of reference genomes available: the *A. thaliana* reference has seven scaffolds with an N50 of 23,459,830, whereas the *A. halleri* reference has 282,453 scaffolds with an N50 of 17,686. To control for this limitation, we again removed contigs < 10KB from our combination reference genome, which helped with run time and memory usage but did not affect the conclusions (Figure S10b). These tests in *Arabidopsis* provide an empirical illustration of sppIDer’s performance with differing quality reference genomes. *Arabidopsis* also provides a useful test of detecting hybrids in a plant system, as there are two well-supported allotetraploid species in the genus, *Arabidopsis suecica* and *Arabidopsis kamchatica* (Shimizu-Inatsugi et al. 2009; Schmickl et al. 2010). First, we tested short-read data from a divergent lineage of *A. thaliana* (Durvasula et al. 2017) and found that the reads mapped well to the *A. thaliana* reference genome (Figure 5d). As expected, reads from the interspecies hybrid *A. kamchatica* (Novikova et al. 2016) mapped both to *A. lyrata* and to *A. halleri* (Figure 5e), approximately equally, confirming that *A. kamchatica* indeed has genomic contributions from these two species and that sppIDer can detect hybrids, even when the combination reference genome contains reference genomes of substantially varying quality. Thus, sppIDer can accurately detect interspecies hybrid in a plant model and will likely become more generally useful in other plant systems, where allopolyploidy is frequent (Soltis et al. 2015), as more reference genomes become available.

### mitoSppIDer

Applications of sppIDer with non-nuclear sequencing data are also of considerable interest. Organelle genomes (e.g. mitochondria, chloroplast) have a different mode of inheritance, and increasing data suggest widespread reticulation and cases where their ancestries differ from the nuclear genomes (Peris et al. 2014; Wu et al. 2015; Leducq et al. 2017; Peris et al. 2017a; Peris et al. 2017c; Sulo et al. 2017). We developed mitoSppIDer as an extension to explore these non-nuclear inherited elements. Since mitochondrial genomes are generally small, the coding regions can be easily visualized, which allows precise mapping of introgressions in both coding and non-coding regions. However, more cautious interpretation is warranted, because mitochondrial reads are often at low and variable abundance, and quality can differ between DNA isolations and sequencing runs. Again, we tested using the genus *Saccharomyces* because of the availability of mitochondrial reference genomes (Foury et al. 1998; Procházka et al. 2012; Baker et al. 2015). We first tested mitoSppIDer with a strain of *S. uvarum* (ZP1021) (Almeida et al. 2014) and found that, of the reads that mapped to any mitochondrial genome, > 99%mapped to the *S. uvarum* mitochondrial genome (Figure S12a). Next, we examined Vin7, a hybrid strain of *S. cerevisiae* X *S. kudriavzevii*, and mitoSppIDer revealed that this strain inherited the mitochondrial genome of *S. kudriavzevii* with intergenic introgressions from multiple non-*S. kudriavzevii* mitochondrial genomes (Figure S12b) (Peris et al. 2017c). As with conventional sppIDer, mitoSppIDer rapidly highlights interesting regions for further analysis, such as detailed phylogenetic analyses of introgression candidates.

### Summary

Altogether, these tests show the versatility of sppIDer across clades: in fungi, plants, and animals. sppIDer allows for the rapid exploration and visualization of short-read sequencing data to answer a variety of questions, including species identification; determination of the genome composition of natural, synthetic, and experimentally evolved interspecies hybrids; and inference of CCNVs (Brickwedde et al. 2017; Gorter de Vries et al. 2017; Peris et al. 2017b). With examples from the genus *Saccharomyces*, sppIDer could detect contributions from up to four species and recapitulated the known relative ploidy and aneuploidies of brewing strains. From a simulated phylogeny, we found that sppIDer accurately detected hybrids from a range of divergences in the parents and even detected ancient hybrids. In systems with low-quality or varying quality references genomes, sppIDer performs well without much promiscuous mapping between varying reference qualities, but its ability to infer translocations and CCNVs is limited. Even in systems missing reference genomes, sppIDer still enables rapid inferences by using the reference genomes of closely related species, with the caveat that mapping quality declines with sequence divergence. Additionally, sppIDer works on long-read data and with coverage as low as 0.5X. Finally, sppIDer can be extended to non-nuclear data, allowing for the exploration of alternative evolutionary trajectories of mitochondria or chloroplasts. As more high-quality reference genomes become available across the tree of life, we expect sppIDer will become an increasingly useful and versatile tool to quickly provide a first-pass summary and intuitive visualization of the genomic makeup in diverse organisms and interspecies hybrids.

## Methods

The sppIDer workflow to identify pure species, interspecies hybrids, and CCNVs consists of one main pipeline that utilizes common bioinformatics programs, as well as several custom summary and visualization scripts (Figure 1). An upstream step is required to prepare the combination reference genome to test the desired comparison species. The inputs for the main sppIDer pipeline are this combination reference genome and short-read FASTQ file(s) from the organism to test. The output consists of several plots showing to which reference genomes the short-reads mapped, how this mapping varies across the combination reference genome, and several text files of summary information. Additionally, the pipeline retains all the intermediate files used to make the plots and summary files; these contain much more detailed information and may be useful as inputs to various other potential downstream analyses. We are releasing sppIDer as a Docker, which runs as an isolated, self-contained package, without the need to download dependencies and change environmental settings. Packaging complex bioinformatics pipelines as Docker containers increases their reusability and reproducibility, while simplifying their ease of use (Boettiger 2015; Di Tommaso et al. 2015). sppIDer can be found here (https://github.com/GLBRC/sppIDer), where a transparent Dockerfile lays out the technical prerequisites, platform, how they work in combination, and is a repository for all the custom scripts. A manual for sppIDer can be found both at the GitHub page and at http://sppider.readthedocs.io.

### The pipeline

Before running the main sppIDer script, a combination reference genome must first be created and properly formatted (top of Figure 1). This is a separate script, combineRefGenomes.py, that takes multiple FASTA-formatted reference genomes and a key listing the reference genomes to use and a unique ID for each. The script concatenates the reference genomes together in the order given in the input key, outputting a combination reference FASTA where the chromosomes/scaffolds are renamed to reflect their reference unique ID and their numerical position within the reference-specific portion of the combination output. For reference genomes that contain many short and uninformative scaffolds, there is an option to remove scaffolds below a desired base-pair length. This option improves speed, memory usage, and visual analysis for large genomes with many scaffolds and low N50 values. Setting a threshold usually does not affect the conclusions (Figure S10), but we recommend trying different thresholds to determine how much information is lost. The choice of reference genomes to concatenate is completely at the discretion of the user and their knowledge of the system to which they are applying sppIDer. We recommend choosing multiple phylogenetically distinct lineages or species, where gene flow and incomplete lineage sorting are limited, from a single genus. We caution that, for ease of analysis and interpretation, less than 30 reference genomes should be used at once. To illustrate the power of sppIDer, for our examples, we used all available species-level reference genomes for the genera tested, but we excluded lineages and strains within species. However, sppIDer could be applied iteratively with different combinations of reference genomes that are more targeted for a particular lineage or question. For example, with an experimentally evolved hybrid, just the parental genomes could be included to detect CCNVs that occurred during the evolution, but with a suspected hybrid isolated from the wild or industry, all potential parent species reference genomes should be included.

The main body of sppIDer (Figure 1b) uses a custom python 2 (Python Software Foundation) script to run published tools and custom scripts to map short-reads to a combination reference genome and parse the output. The first step uses the mem algorithm in BWA (Li and Durbin 2009) to map the reads to the combined concatenated reference genome. Two custom scripts use this output to count and collect the distribution of mapping qualities (MQ) for the reads that map to each reference genome and produce plots of percentage and MQ of reads that map to each reference genome. The BWA output is also used by samtools view and sort (Li et al. 2009) to keep only reads that map with a MQ > 3, a filter that removes reads that map ambiguously. From here, the number of reads that map to each base pair can be analyzed using bedtools genomeCoverageBed (Quinlan and Hall 2010), for smaller genomes using the per-basepair option (-d) and, for large genomes, the –bga option. The depth of coverage output is used by an R (Wickham 2009; R Core Team 2013) script that determines the mean coverage of the combined reference genome that is subdivided into 10,000 windows of equal size. Finally, a plot for the average coverage for each component reference genome and a second plot of average coverage for the windows are produced.

### The metrics

Several different metrics are used to summarize the data. Depth of coverage is a count of how many reads cover each base pair or region of the genome. Coverage can vary greatly from sequencing run to sequencing run; hence, a log_2_ conversion is used to normalize to the mean coverage. As discussed in the Results, depth of coverage plots can be used to infer the species, the parents of hybrids, and ploidy changes either between or within a genome. sppIDer also reports the percentage of reads that map to each reference genome. Finally, sppIDer uses the established MAPPing Quality (MQ) score introduced in Li et al. (2008) to bin reads by their map-ability on a 0-60 scale. A score of zero is used for reads where it is unlikely that their placement is correct, so sppIDer reports these as “unmapped”, along with reads that cannot be mapped and therefore do not receive a MQ score. The mapping quality scale can therefore provide a rough assessment of data quality, as well as divergence to the provided reference genomes.

### Tested reference genomes and data

For the *Saccharomyces* tests, we used reference genomes that are scaffolded to a chromosomal level. In some cases, there is only one reference genome available per species, and for the others, we used the first available near-complete reference; see Table S1 for those used. For systems with multiple reference genomes available, the choice could be more targeted, such as utilizing lineage specific references or references that contains unplaced scaffolds with genes of interest. Alternatively, for systems where few genomes are available, we have shown here that a close relative works as a proxy. For the *Saccharomyces* references, each ordered “ultra-scaffolds” genome was downloaded from http://www.saccharomycessensustricto.org/ or for *S. arboricola* and *S. eubayanus* from NCBI. The published *S. uvarum* genome (Scannell and Zill et al 2011) had chromosome X swapped with chromosome XII, which was fixed manually. These genomes were concatenated together using the python script combineRefGenomes.py, creating a combination reference FASTA with all *Saccharomyces* species. This combination reference genome can then be used repeatedly to test any dataset of interest. For the *Saccharomyces* tests, we used publicly available FASTQ data from a number of publications, all available on NCBI (Table S1 contains all accession numbers). Using the data for each strain separately and the combination reference genome created above, we then called sppIDer.py with, --out uniqueID, -- ref SaccharomcyesCombo.fasta, --r1 read1.fastq, and optionally --r2 read2.fastq. sppIDer is written to test one sample’s FASTQ file(s) against one combination reference genome at a time, but this could be easily parallelized.

For the tests to determine if hybrids could be detected with missing reference genomes, new combination reference genomes without one species’ genomes were created by removing the desired species’ reference name from the reference genome key before running combineRefGenomes.py. Since both Vin7 and W34/70 contain contributions from *S. cerevisiae,* the combination reference genome lacking the *S. cerevisiae* reference was tested for each set for FASTQ files for Vin7 and W34/70. The same process was followed to remove the *S. kudriavzevii* reference genome from the combination reference to test Vin7, as well as to remove the *S. eubayanus* reference genome from the combination reference to test W34/70.

For the *Lachancea* test, all of the genomes were available and downloaded from http://gryc.inra.fr/. The FASTQ data for CBS6924 was downloaded from NCBI. A combination reference genome with all available genomes was created and used. Then, sequentially, the “*Lachancea fantastica*” and *Lachancea lanzarotensis* genomes were removed by modifying the input key and rerunning combineRefGenomes.py. The FASTQ data for CBS6924 was tested against all three of these combination reference genomes. See Table S1 for the full accessions.

For the non-*Saccharomyces* tests, we used the most complete reference genome available for each species in the genus (accessions Table S1). Therefore, there is quite a bit of variation between different references. For the *Drosophila* and *Arabidopsis* genomes, we tested removing contigs, using the --trim option, with combineRefGenomes.py, as well as not removing contigs, and found the cleanest results when we removed contigs less than 10 Kb. The combined reference genomes of both *Drosophila* and *Arabidopsis* were both larger than four gigabases; therefore, the --byGroup option was used with sppIDer.py to speed up processing and reduce memory usage. The data we tested came from a variety of publications, but we targeted data of divergent or hybrid lineages. See Table S1 for complete information.

For the mitoSppIDer test, we used the complete species-level *Saccharomyces* mitochondrial reference genomes available on NCBI, which do not necessarily correspond to the same strain that was used to build nuclear genomic reference (Table S1). Again, combineRefGenomes.py was used to concatenate these references. An additional script, combineGFF.py, was used to create a combination GFF file that was used to denote the coding regions on the output plots. mitoSppIDer.py has an additional flag for the GFF file, but it otherwise runs in a similar manner to sppIDer.py; the same input FASTQ file(s) can even be used. Whole genome sequencing data contains varying amounts of mitochondrial sequences; therefore, using the raw FASTQ data works sufficiently, even when many of the genomic reads will be classified as “unmapped”.

### Simulations

To create the simulated low-quality de novo genomes, we used the software iWGS (Zhou et al. 2016) to simulate 100bp paired-end reads with an average inter-read insert size of 350bp (sd 10) at 2X coverage from the reference genomes of *S. cerevisiae, S. kudriavzevii, S. uvarum,* and *S. eubayanus*. For the simulated de novo *Saccharomyces* genomes the N50 scores ranged from 1254-1274 and the number of scaffolds ranged from 10023-10426 (Table S1).

To simulate short-read data, we used DWGSIM (https://github.com/nh13/DWGSIM), which allowed us to vary the coverage, error rate, and mutation rate as needed. The *S. cerevisiae* reference genome was used to simulate single species reads and a concatenation of the *S. cerevisiae* and *S. eubayanus* reference genomes was used for hybrid pseudo-lager reads. As a test of an aneuploid genome, we also manually manipulated the *S. cerevisiae* reference genome so that it contained zero copies of chromosomes I and III and duplicate copies of chromosome XII,. All simulated reads were 100bp paired-end reads with an average insert size of 500bp. For the coverage tests, we varied the coverage from 0.01-10X. For the short reads used against the low-quality de novo genomes, we used 10X coverage and a 3% mutation rate. To simulate PacBio-style long reads, we used iWGS on the hybrid pseudo-lager concatenated genome with the default settings of 30X coverage, average read accuracy of 0.9, and SD of read accuracy 0.1.

To make our simulated phylogeny, we used the *S. cerevisiae* reference genome as a base and simulated reads with DWGSIM at a 2%mutation rate as 100bp paired-end reads with an average insert size of 500bp at 10X coverage. iWGS was used to assemble these reads. The resulting assembly was again simulated with a 2% mutation rate, and those reads were assembled. This procedure was followed for 6 rounds with one lineage being independently simulated twice each round to produce a speciation event. This simulation resulted in 10 species in the phylogenetic arrangement shown in Figure 3a. Summaries of the final assemblies can be found in Table S1, but the median of the final assemblies was 5100 scaffolds, N50 of 1335, and total length of 6.4MB. Each simulated species was ∼12% diverged from *S. cerevisiae,* the most closely related species were ∼4% diverged, and the most distantly related species were ∼20% diverged. The reads used to produce the final assemblies were used to test whether sppIDer mapped each set of reads to their corresponding reference genomes. The reads of different references were concatenated to simulate pseudo-hybrids of different divergences. To simulate ancient hybrids, the reads from earlier rounds of simulation, before speciation events, were concatenated and tested against the final assemblies with sppIDer. As with the empirical data, to simulate a missing reference genome, that reference was removed from the input key prior to running combineRefGenomes.py.

### Alignment-free phylogenetic methods

We tested four alignment-free phylogenetic methods: two that build phylogenies using short-read data, SISRS (Schwartz et al. 2015) and AAF (Fan et al. 2015), and two that assemble targeted loci from short-read data, aTRAM (Allen et al. 2015) and HybPiper (Johnson et al.2016). We simulated 10X coverage paired-end, 100bp data for each *Saccharomyces* reference genome at a mutation rate of 0 with DWGSIM to use as input for these methods. For SISRS, we used the default settings with a genome size of 12Mb, first using only the reference *Saccharomyces* data, then including empirical data for hybrids. SISRS failed at the missing data filtering step when data from the lager strain W34/70 was used, even when we allowed for all but one sample to have missing data. SISRS nexus outputs were visualized with SplitsTree (Huson and Bryant 2006). For AAF, we found that a *k* of 17 accurately recapitulated the *Saccharomyces* phylogeny, even with the inclusion of empirical data from other pure lineages. Once we determined the optimal *k*, we tested including empirical hybrid data. We also used AAF with our simulated phylogeny, which constructed the tree that matched the simulations with the default *k* of 25. The output of AAF was visualized with iTol (Letunic and Bork 2016).

For the targeted loci methods, we used 13 loci that can delineate *S. eubayanus* populations (Peris and Langdon et al. 2016), as well as the ITS sequences for *S. cerevisiae* (AY046146.1) (Kurtzman and Robnett 2003) and *S. eubayanus* (JF786673.1) (Libkind and Hittinger et al. 2011) as bait, all obtained from NCBI. We tested the simulated *Saccharomyces* reads, as well as the empirical data for P1C1, Fosters O, CBS1503, CBS2834, Vin7, and W34/70. For aTRAM, we used the default settings and the option for the Velvet assembler. For HybPiper, we used the default settings and the SPADES assembler.

## Acknowledgments

We thank Drew Doering for the portmanteau sppIDer and EmilyClare Baker and María Lairón Peris for beta testing. This material is based upon work supported by the National Science Foundation under Grant Nos. DGE-1256259 (Graduate Research Fellowship to QKL) and DEB- 1253634 (to CTH), the Robert Draper Technology Innovation Fund from the Wisconsin Alumni Research Foundation (to CTH), the USDA National Institute of Food and Agriculture under Hatch Project 1003258 (to CTH), and funded in part by the DOE Great Lakes Bioenergy Research Center (DOE BER Office of Science DE-SC0018409 and DE-FC02-07ER64494). QKL was also supported by the Predoctoral Training Program in Genetics, funded by the National Institutes of Health (5T32GM007133). DP is a Marie Sklodowska-Curie fellow of the European Union’s Horizon 2020 research and innovation programme, grant agreement No.747775. CTH is a Pew Scholar in the Biomedical Sciences and a Vilas Faculty Early Career Investigator, supported by the Pew Charitable Trusts and the Vilas Trust Estate, respectively.

